# In vivo imaging defines vascular interplay in the development of lymphocytic leukemia in zebrafish models

**DOI:** 10.1101/806562

**Authors:** Sergei Revskoy, Margaret E. Blair, Shaw M. Powell, Elizabeth S. Hausman, Jessica S. Blackburn

## Abstract

The vascular microenvironment at the primary tumor site is represented by functionally diverse types of vessels that contribute to metabolic supply, trafficking/dissemination of tumor cells, and delivery of chemotherapeutics. However, the role of leukemia-associated vascular networks in leukemia progression remains poorly understood. We utilized a MYC-induced T cell leukemia model in zebrafish to assess the involvement of vascular endothelial growth factor A-dependent (VEGFR2+), VEGFA-independent vasculature (VEGFR2-), and lymphatics in the initial stages of leukemia cell dissemination using intravital microscopy. Leukemogenesis underwent sequential progression, from initial successive emigration of single leukemic cells from the thymus towards the kidney marrow, followed by collective migration in a dorsal-lateral direction with scattered migration from the ventral thymus towards the aortic arches, before streaming towards the rostral kidney area. Trafficking appeared independent of VEGFR2+ vasculature but was closely associated with VEGFR2-microvessels, and, interestingly, leukemic cells did not utilize lymphatics during initial dissemination. Overall, leukemic cells appeared to utilize the same routes as normal T cell progenitors during development but in a reverse order, and primarily recruited VEGFR2-microvessels during the initial stages of progression. Ultimately, interfering with leukemia cell migration by targeting vascular networks may represent a new therapeutic strategy to control leukemia progression.

## 1. Introduction

Immature lymphocytes are a common target for malignant transformation into leukemia. For example, acquisition of high expression of the Myc oncogene is closely involved in lymphocytic leukemia in mammals and zebrafish [1-3]. In addition, while differentiation of recombination activating gene 2 (Rag2)-expressing lymphocyte precursors constitutes a critical step in the development of immunocompetency, the intensive V(D)J recombination can result in Rag2-induced off-target effects that may drive diverse leukemogenesis [4-6]. These types of genetic events might significantly modify and diversify the behavior of the MYC-transformed leukemia cells, and the molecular events accompanying Myc-induced cell transformation in leukemia and other cancers have been extensively delineated. However, the microenvironmental and developmental contexts of leukemogenesis remain largely unknown.

Normal Rag2+ T cell progenitors that colonize the thymus undergo a well-established life cycle, including migration into the primordial thymus, homing to specific compartments within thymus, and either apoptosis or clonal expansion followed by further lineage-directed differentiation and emigration away from the thymus. In T cell leukemia, with high Myc expression in particular, normal Rag2-expressing lymphocyte differentiation is blocked and cells are prompted towards uncontrolled proliferation [7-9]. In addition to intrinsic changes in the transformed cells, Myc overexpression in other types of cancers causes inflammatory remodeling of the tumor microenvironment, angiogenesis, and ultimately, promotes metastatic behavior [10,11]. Unlike the metastatic spread of epithelial cancers, lymphocytic leukemia dissemination is generally not a reflection of tumor progression, but rather of a conserved physiological behavior that might, at least partially, recapitulate trafficking of normal lymphocytes and their progenitors. However, the routes that normal and leukemic lymphocytes utilize for trafficking in and out the thymus are not well-defined [12,13]. Even less is known about the roles of different vascular networks in the thymus; the vascular contribution to metabolic supply of the thymus may ultimately affect cell behavior.

The conventional view of leukemic dissemination is that the growing clones extend through contiguous interstitial spaces around the thymus due to increased physical pressure on the surrounding tissue by the accumulating cell mass [14]. This notion implies that the microenvironment surrounding the thymus does not have a significant role in leukemia dissemination during the initial stages of the process. Alternatively, it can be speculated that lymphatic neoplasia that originates from more differentiated T cells might conserve lymphatic cell migration behavior, and disseminate in a less chaotic manner, i.e. gradually emigrating from thymus via efferent vessels or other routes from the exponentially growing colony *in situ*. Additionally, as seen in solid tumors, the leukemic cells themselves may remodel the vascular microenvironment to accommodate intra- and extravasation, and changing metabolic needs of the growing leukemia. A more complete understanding of routes of dissemination and leukemic cell interaction with the microenvironment are important issues in leukemogenesis that remain unresolved. Additionally, the spatiotemporal aspects of tumor angiogenesis have only marginally been explored *in vivo*, and only in solid tumors.

Zebrafish models provide valuable insight into both cancer progression and angiogenesis. The morphology, molecular mechanisms of induction, biological behavior, and response to treatment of malignant tumors in zebrafish are very similar to those in mammals. Indeed, most of the tumors induced in zebrafish by either carcinogens or oncogenic transgenes behave similarly to mammalian cancers with uncontrolled proliferation, loss of differentiation, aneuploidy, and metastases [15]. These features are also characteristic of zebrafish models of leukemia caused by forced expression of transgenic oncogenes like Myc in lymphocyte progenitor cells [3,16]. Angiogenesis in zebrafish is also similar to that of mammals, and has been well characterized by using VEGFR2 (Kdlr) and Fli1 transgenic reporter systems [17-19]. Tumor-related angiogenesis has also been demonstrated in zebrafish, in both early embryos and in xenograft systems in adult fish [20-22], although the latter models are profoundly immunosuppressed and may not provide an accurate representation of the tumor-vasculature crosstalk. Importantly, the interactions between tumor and vasculature has not been addressed in a context of developing primary tumor *in vivo*.

Here, we have begun to address these issues by visualizing the process of leukemogenesis *in vivo* using the zebrafish model of T cell lymphocytic leukemia (a *rag2:GFP-Myc* transgenic fish line), documenting chronologically distinct steps in tumor development, and observing leukemic cell interactions with various types of vasculature, including Vascular Endothelial Growth Factor Receptor (VEFGR2)+ and VEGFR2-endothelial vessels and lymphatics. By establishing the pattern of leukemia cell migration and their routes, we will be able to decipher mechanisms of leukemia dissemination and, ultimately, control this process. In addition, establishing the vascular microenvironment present in the early stages of leukemogenesis would provide better insight into mechanisms of metabolic reprogramming that is needed during leukemic transformation and spread. Given the heterogeneous nature of leukemia, with lymphocytic leukemias having several distinct clinical subtypes linked with patient prognosis, and leukemia cells within even the same patient harboring genetically distinct clones, defining the role of microenvironmental factors in leukemia progression may provide novel therapeutic strategies to more broadly control leukemia dissemination and progression.

## 2. Results

### 2.1 Visualization of step-wise leukemia onset in vivo in a zebrafish lymphocytic leukemia model

To obtain lymphocytic leukemia in zebrafish, we utilized a well-established model for Myc-induced thymic-derived leukemia [16]. In a sub-line derived from this *rag2:GFP-Myc* model, the developing leukemia is restricted to the cortical part of the thymus, which is the primary site of Rag2+ cells, and demonstrates high penetrance, early onset, progression towards dissemination in a stepwise manner, and a relatively benign course of the disease, so that the leukemia-bearing fish remain fertile [23].

We outcrossed these fish, as well as a *Rag2:GFP* control line, to the Casper stain [24] for ease of imaging using Light Sheet microscopy. Interestingly, the *Rag2:GFP-Myc* offspring consistently demonstrated the same pattern of leukemia progression, including almost synchronous onset, even in heterozygous animals. At 11-14 days post-fertilization (dpf), we observed (n>30 animals for all observations) that Rag2:GFP and Rag2:GFP-Myc expressing cells in thymus were confined to a pouch-shaped cortical area and demonstrated a similar growth pattern whether they expressed Myc or not (Figure 1 A, B). Beginning at 12-14dpf, thymic growth of Myc+ fish significantly accelerated into a thymic hyperplasia, and single Rag2:GFP-Myc expressing leukemic cells began to depart from the caudal tip of the organ in a uni-directional manner, resembling a channeled trafficking that appeared to follow a curvature directed internally towards the kidney, which is the site of the marrow in zebrafish (Figure 1C). This was followed by a scattered migration in a ventral direction (Figure 1C), then, around 21dpf, a massive dorsal-lateral emigration began towards the medullar thymus and the vascular plexus associated with the kidney (Figure 1D).

**Figure 1.**
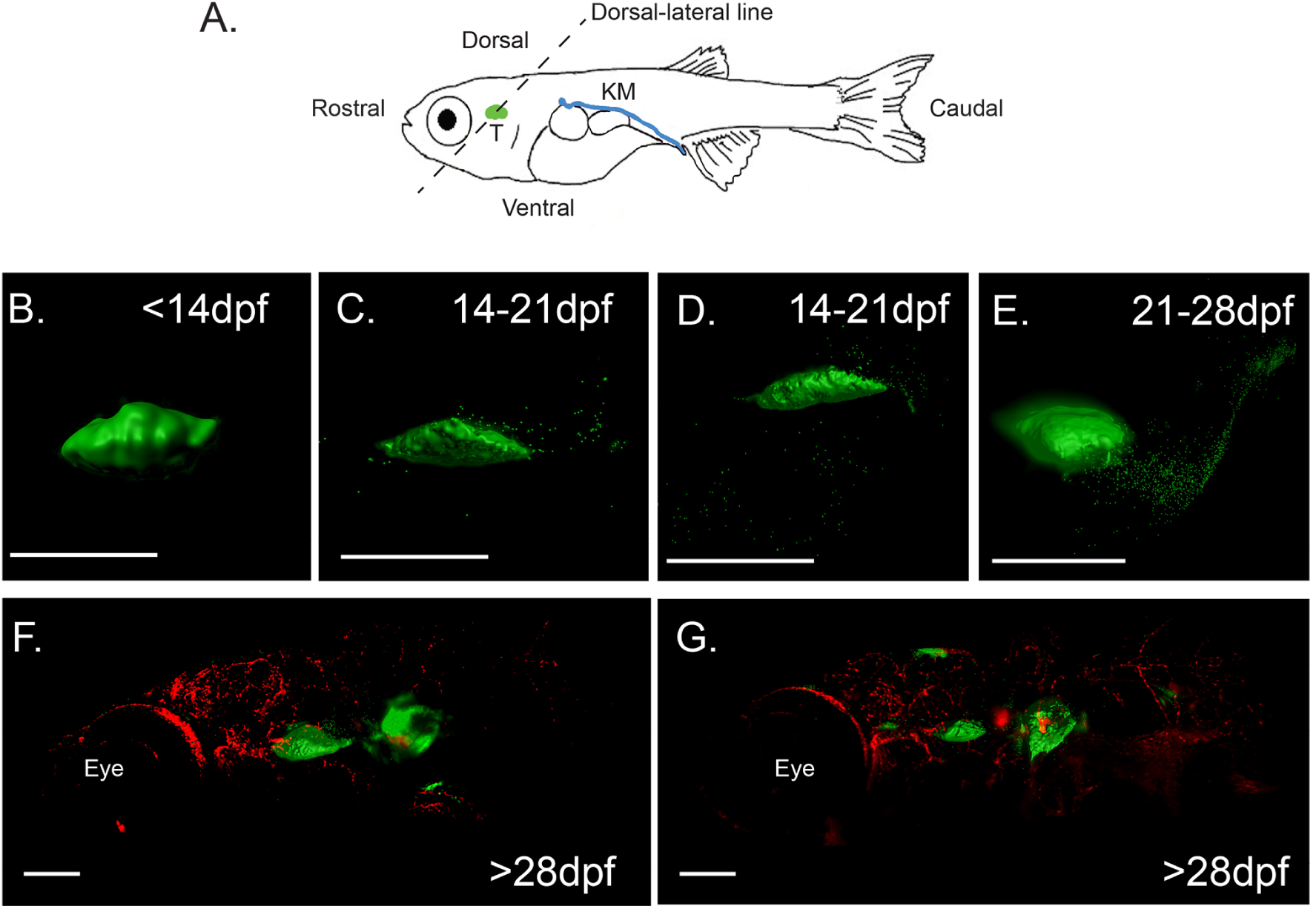
Lymphocytic leukemia in *Rag2:GFP-Myc* fish progresses in a step-wise and time dependent manner. (**a**) Schematic of zebrafish with location of thymus (T) and kidney marrow (KM) shown, and terms that define directionality in the animal are noted. Light sheet microscopy was used to image the thymus and GFP-Myc+ leukemia cells to show leukemia progression. At <14dpf (**a**) cells remain in the thymus, but start to move out from the caudal tip (**c**) as well as ventrally (**d**) at 14-21dpf. At 21-28dpf (**e**) cells begin a mass emigration to the kidney marrow. At late stages, >28dpf, there are large leukemic colonies in the kidney marrow (**f**) and other sites in the body (**g**). Scale bar = 50μm in (**b-e**), and 200μm in (**f-g**). Flk:RFP is shown in (**f**-**g**) to orient the whole body image.

After 28 dpf, cells quickly spread into kidney head before gradually expanding to kidney trunk and developing large colonies there, followed by movement into the area corresponding to the caudal vein plexus before dissemination throughout the body (Figure 1E). At this stage of leukemia progression, leukemia colonies also emerged in the posterior cranial area, a prospective area of adult-like kidney and in the retro-orbital area, near the former pharyngeal patch, the site which gave rise to thymic epithelium (Figure 1F). Why subsets of leukemic cells prefer to home to one niche over others is an area that we are actively investigating.

In total, these data show that leukemic cells are not emigrating from the thymus and disseminating in a stochastic manner. Instead, leukemogenesis appears to happen in a sequential and directional manner that was consistent in all animals that we observed. Myc-derived zebrafish T cell leukemias are known to have heterogenous mutations and differential gene expression profiles, between individual leukemias and between clones within the same leukemia [14,25,26]. Yet, our data that leukemias consistently follow the same dissemination pattern suggest that initial leukemia onset is not reliant on acquisition of certain mutations or gene expression, and instead may be heavily influenced by either normal routes of T cell emigration from the thymus and/or external factors such as the thymic microenvironment.

### 2.2 VEGFR2+ vasculature is not spatially associated with Rag2:GFP-Myc leukemia dissemination

The vascular microenvironment is key to cancer progression as it influences the metabolic needs of the tumor and provides a route for solid tumor cell dissemination. The contributions of the vascular system to leukemogenesis are unknown. To gain insight into how the vascular microenvironment might impact the initial steps of dissemination of a developing leukemia in the thymus, we crossed *rag2:GFP-Myc* transgenic fish with lines of fish harboring transgenic reporters that label different types of vasculature, including *flk:RFP*, which labels all VEGFR2+ vasculature [27,28], *lyve:dsRED*, which labels lymphatics [29], and *fli1ep:GAL4FF/UAS:mRFP*, which is based on the pan-vascular Fli1a but is negatively regulated by VEGF-A [19,30,31], and which in our model depicts VEGFR2 negative microvasculature.

VEGF-A, the main vascular growth factor, acts primarily via VEGFR2 receptor on endothelial cells, promoting arterial-venous angiogenesis and vascular integrity in various tissues and organs under physiologic conditions. It is well documented in a variety of cancers that VEGFA secretion by tumor cells promotes angiogenesis in VEGFR2+ endothelium, and this process is thought to be critical for growing tumors to remain oxygenated. Interestingly, we observed no role for VEGFR2+ vasculature in T cell leukemia onset. By 14dpf, VEGFR2+ main vascular branches and their ramifications were not spatially associated with normal Rag2:GFP thymus and even to a lesser extent with leukemic ones (Figure 2, and quantified in Figure 3A). The oxygen-rich VEGFR2+ vasculature was primarily present in aortic arches, which are ventral to the cortical thymus. Surprisingly, by 21dpf, only sparse individual Rag2:GFP-Myc leukemic cells were associated with this VEGFR2+ vasculature, but cells neither embedded nor intravasated into these vessels until wide-spread dissemination of leukemia at later stages (>28dpf) of leukemia progression (data not shown). As Rag2:GFP-Myc leukemia progressed to 28dpf, VEGFR2+ vasculature remained sparse in ventral area of the thymus and barely present in dorsal area, and was actually located a further distance from the leukemic thymus than normal thymus (Figure 3B, p=0.044). At 28dpf, massive collective emigration of Rag:GFP-Myc leukemic cells from dorsal lateral area of thymus and continuous emigration of cells from the caudal tip of the thymus towards the kidney/marrow occurred essentially in VEGFR2-free zone (Figure 2).

**Figure 2.**
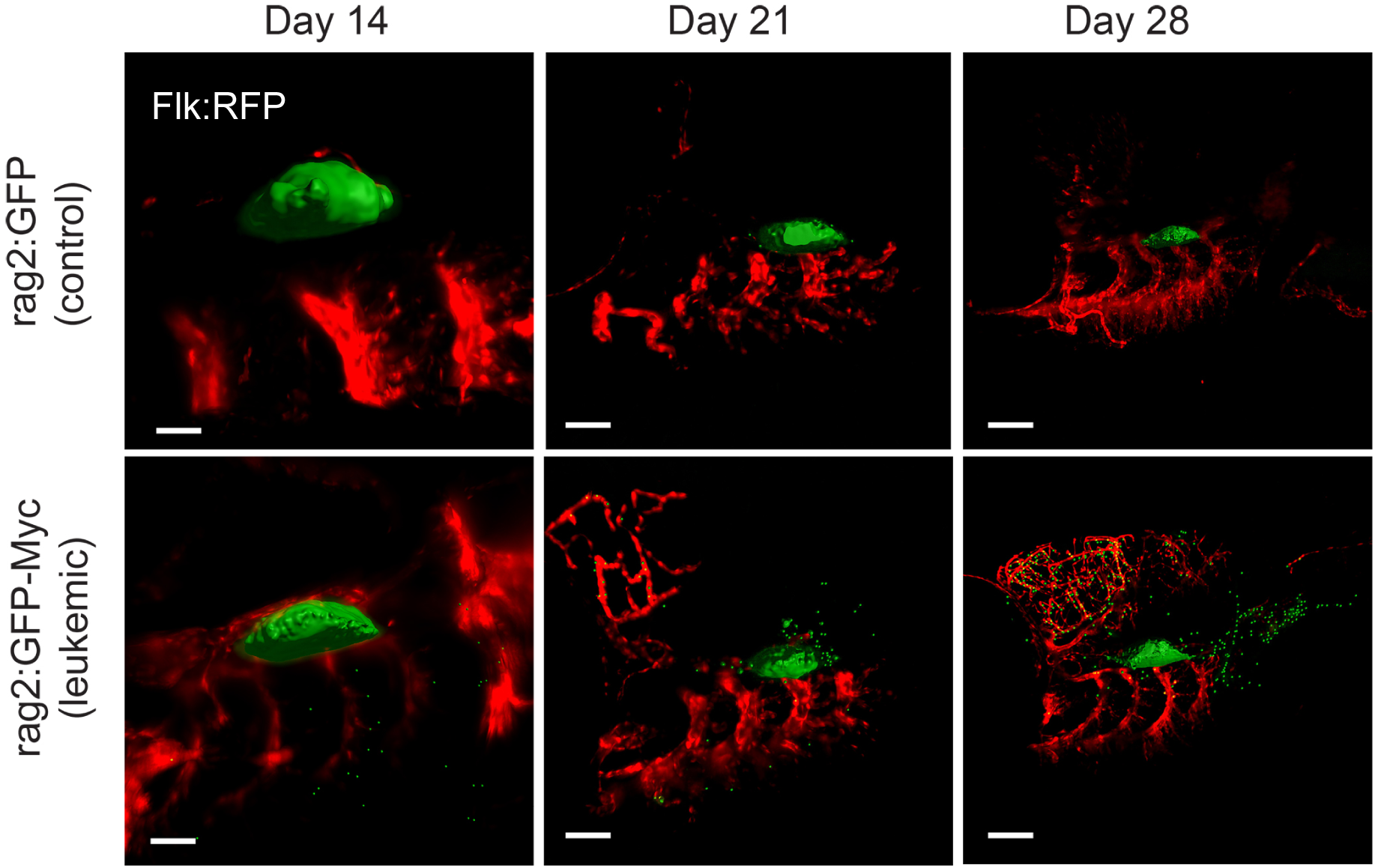
VEGFR2+ vasculature is not associated with leukemogenesis. Flk:RFP;Rag2:GFP and Flk:RFP;Rag2:GFP-Myc zebrafish were examined at 14, 21, and 28dpf, and the thymic area was imaged by light sheet microscopy. VEGFR2+ vasculature was not in close proximity to the thymus at any stage. Subsets of GFP-Myc cells would migrate ventrally into VEGFR2+ aortic arches (14 and 21dpf) but we did not observe close association or intravasation between leukemic cells and VEGFR2+ vessels at this site. At later stages (28dpf), mass doral-lateral migration of leukemia cells towards the kidney occurred in areas that appeared free of VEGFR2+ vasculature. Scale bar=50μm at 14dpf and 150μm at 21 and 28dpf.

**Figure 3.**
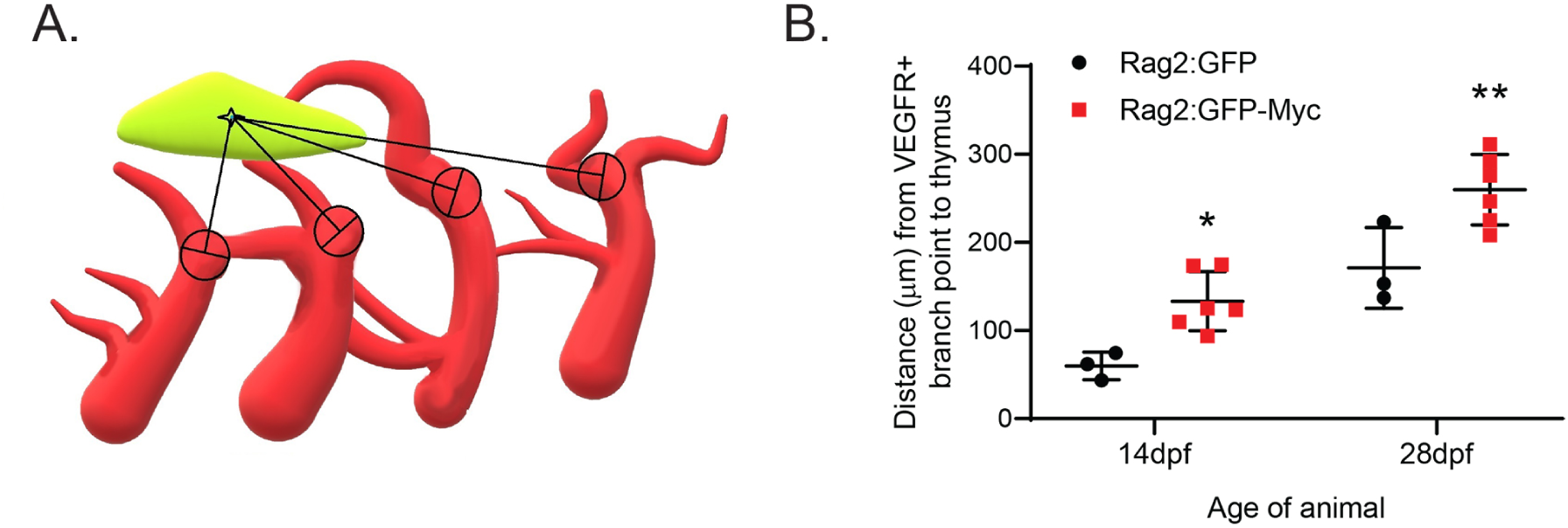
VEFGR2+ vasculature is not in close proximity to the leukemic thymus. (**a**) Schematic of measurements, where the distance between the closest branch point of the VEGFR2+ vessels and the thymus was quantified using Imaris 9.2. (**b**) VEGFR2+ vasculature is significantly more distant from leukemic thymuc (Rag2:GFP-Myc) than normal thymus (Rag2:GFP) at both early (14dpf) and late (28dpf) stages of leukemia onset. * *p*=0.001 and \*\**p*=0.044

### 2.3 Lymphatic vasculature is associated with progression of Rag2+ leukemia

Lymphatic vessel branches derived from the facial lymphatics were associated with the dorsal part of the thymus, along the rostral-caudal axis. We did not observe a clear association of lymphatic vessels with the route utilized by single cell emigration via the caudal tip of the thymus between 21-28dpf (Figure 4). However, by 28dpf, we consistently observed a subset of disseminating leukemic cells from dorsal lateral thymus that surrounded the lymphatic vessel and would occasionally intravasate it (Figure 5A).

**Figure 4.**
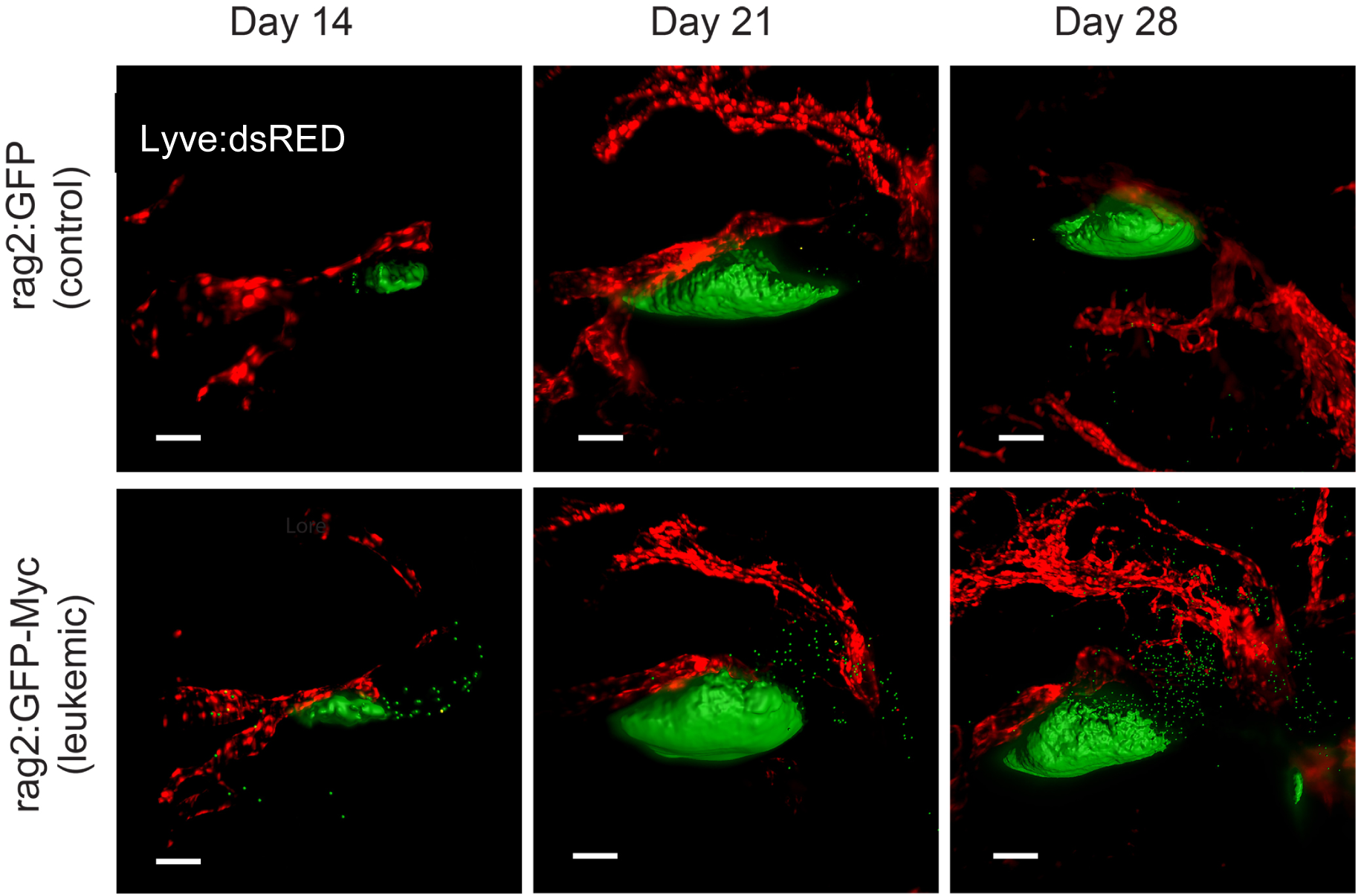
Lymphatics plays a minimal role in leukemogenesis. Lyve:dsRED;Rag2:GFP and Lyve:dsRED;Rag2:GFP-Myc zebrafish were examined at 14, 21, and 28dpf, and the thymic area was imaged by light sheet microscopy. Lymphatics were closely associated with the thymus, and leukemic cells were observed intravasating these vessels (14dpf). However, mass emigration of Rag2:GFP-Myc leukemia cells was not closely associated with lymphatic vessels (28dpf). Scale bar=50μm

**Figure 5.**
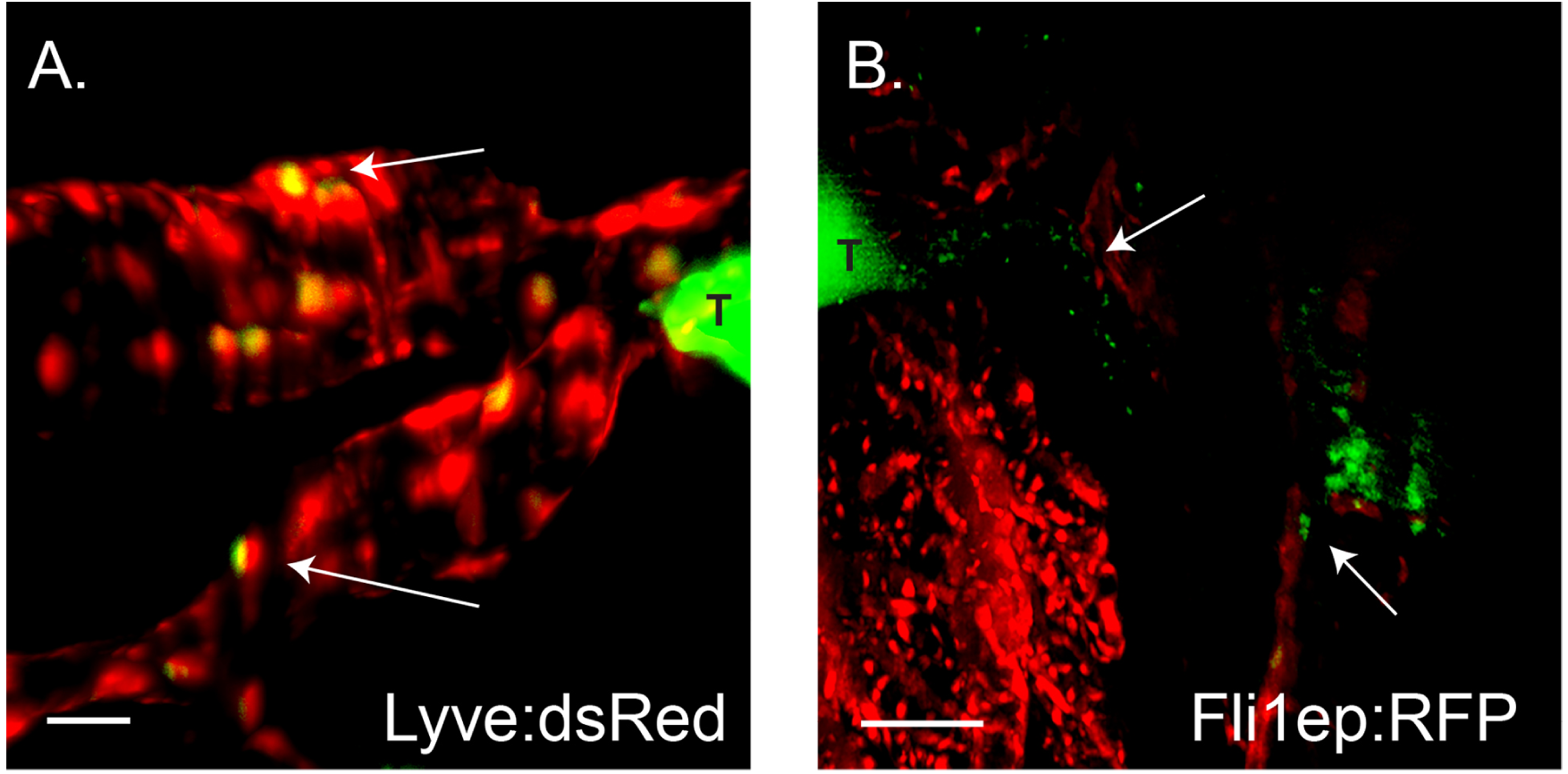
Rag2:GFP-Myc cells are closely associated with lymphatics and VEGFR2-negative endothelial vessels. (**a**) GFP-Myc+ cells can be seen intravasated into Lyve:dsRED lymphatics. (**b**) Leukemia cells appear to line up along the Fli1ep:RFP vessel tracks as the cells emigrate to the kidney marrow. Arrows highlight regions of interest, and T denotes the thymus. Scale bar=15μm in (**a**) and 50μm in (**b**)

### 2.4. Fli1a+ VEGFR2- microvessels vascularize thymus and guide early onset emigration of single leukemic cells

Although all VEGFR-positive vasculature is also positive for Fli1a, a pan-endothelial marker, some Fli1a vessels are VEGFR-negative. We consistently observed that Fli1a+/VEGFR2-negative microvessels (<2µm diameter) penetrated both the normal and leukemic thymus (Figure 6); however, size of the lumen of these vessels prevented them from transporting intravasated leukemic cells, which are approximately 10µm diameter. Instead, these vessels appeared to provide a scaffold for individual leukemic cell migration towards kidney/marrow in early stages of leukemia cell dissemination. We also did not observe any flow of leukemic cells in the larger Fli1a+ vessels surrounding thymus; however, leukemic cells consistently lined up along the outer surface of these vessels, suggesting that they may be crawling along these structures before reaching a secondary site of expansion. (Figure 5B).

**Figure 6.**
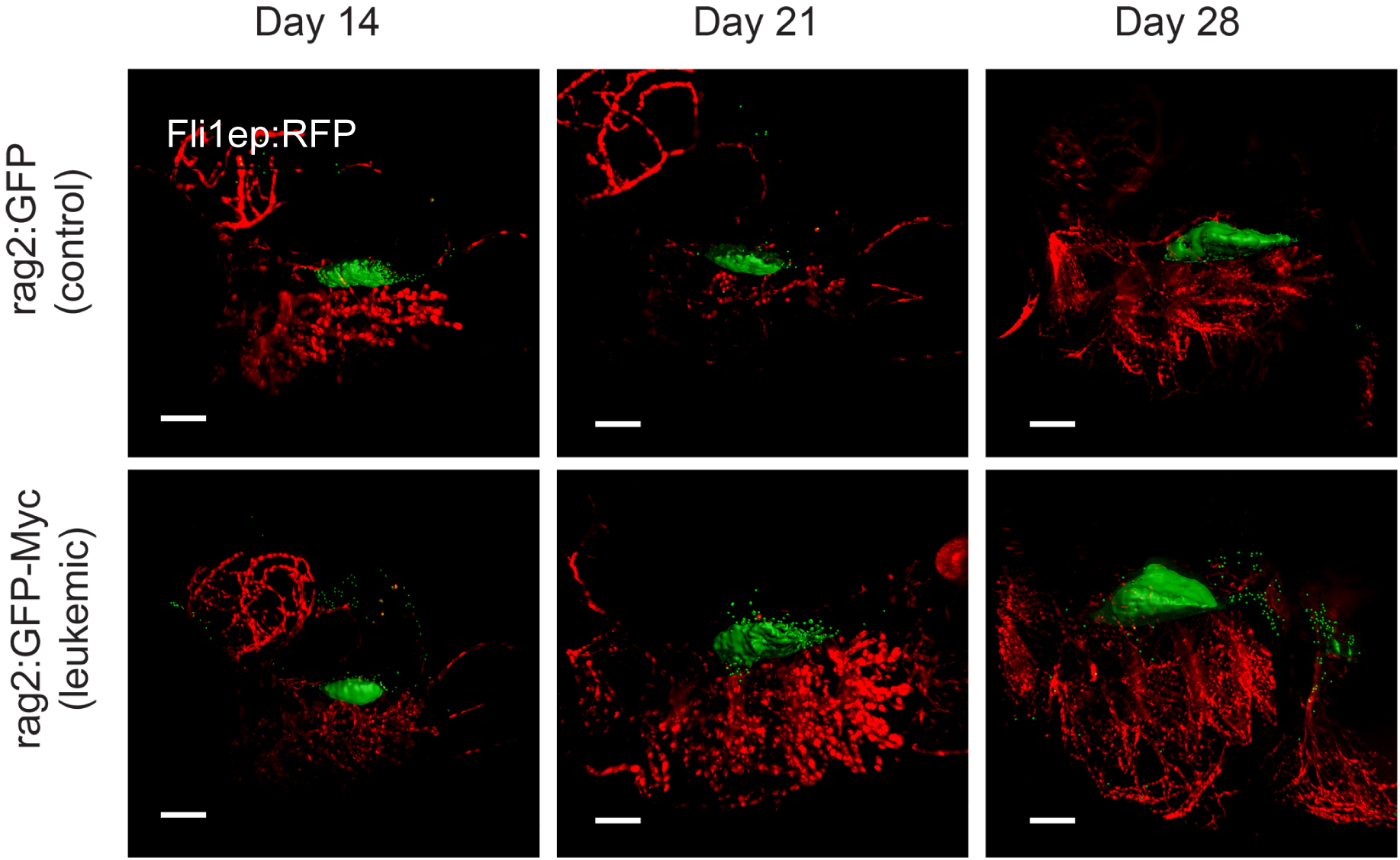
Fli1+/VEGFR2- vasculature and microvessels were closely associated with the thymus and emigrating to the kidney marrow. Fli1epRFP;Rag2:GFP and Fli1ep:RFP;Rag2:GFP-Myc zebrafish were examined at 14, 21, and 28dpf, and the thymic area was imaged by light sheet microscopy. Microvessels can be seen within the thymus at all stages in both leukemic and control animals. At later stages (28dpf), leuekemic cells appear to closely associate with Fli1+/VEGFR2-vasculature as they migrated from the thymus to the kidney.). Scale bar=150μm

In total, we observed leukemogenesis proceeding in a step-wise manner that was reminiscent of normal T cell population of the thymus, but in a reverse order. We found a surprising lack of association of oxygen-rich VEGFR+ vasculature with leukemia dissemination, and limited involvement of lymphatics. Yet, an important role of the vascular component in leukemia onset was apparent in the fli1a+/VEGFR-vasculature, as cells routinely followed trails established by these microvessels. Work to define the specific type of vessel comprised by this population, and it’s precise role in leukemia dissemination, is ongoing.

## 3. Discussion

In this study, we analyzed the pattern of dissemination of Rag2+ leukemic cells in a heritable model of Myc-induced lymphocytic leukemia in zebrafish. Earlier described models of similarly induced zebrafish thymus-derived leukemias showed an acute lymphoblastic leukemia with stochastic diffusion of emigrating leukemic cells, followed by dissemination via blood vessels, with the animal rapidly succumbing to disease [14,16]. In our model, the sequential pattern of leukemic cell dissemination from thymus to marrow to blood, the slow disease progression, and a relatively benign course of leukemia is suggestive of chronic lymphocytic leukemia in mammals, and/or transformation of T cells in later stages of development, and we are in the process of confirming this at the molecular level.

We found that leukemia cell movement away from the thymus largely recapitulates the timing and direction of lymphocyte precursor migration into the primordial thymus, but in a reverse order. Normal lymphocyte precursors utilize multiple routes to reach the thymus in zebrafish, beginning in the narrow region between the dorsal aorta and caudal vein, followed by the pharyngeal arches, the primary head sinus, and the posterior cerebral vein [13,32,33]. In our model of T cell leukemia, Rag2:GFP-Myc cells begin dissemination by leaving the caudal tip of thymic cortex towards kidney area, followed by emigration towards aortic arches, the dorsal lateral area of thymus, and the retro-orbital space. At >28dpf, during later stages of dissemination (data not shown), large number of leukemic cells do enter the blood circulation; however, their gates of entry remain to be established. Overall, the consistent timing of leukemia onset and the similar dissemination route across all animals examined, suggested that, despite potentially unique transforming events, inherent T cell migratory properties and/or the leukemia microenvironment play important roles in regulating leukemia onset.

Additionally, we found that local dissemination of leukemic cells from the thymus sharply declines as distant leukemic infiltrates develop in the area corresponding to the kidney. This raises interesting questions about a crosstalk between secreted migratory factors, which can determine the timing, path, pace, and directionality of leukemic cell migration. Numerous guidance molecules and chemoattractants may specify cells to proximal and distant locations [13,34]; correspondingly, leukemic cells migrating within a lymphoid tissue and between tissues may encounter multiple chemoattractant signals in complex spatial and temporal patterns. Therefore, the interplay between leukemic cells and their microenvironment, including vascular networks, may also change over the course of leukemia progression.

We also observed a consistent spatiotemporal pattern of leukemic cells emigrating from the thymus that was closely associated with vascular context. At early stages of dissemination, Rag2+ leukemic cells demonstrated a lack of any affinity and possibly a repulsive behavior towards VEGF-dependent VEGFR2+ vessels. Although VEGFR+ vasculature plays critical roles in solid tumor progression, the lack of association of VEGFR2+ blood vessels with leukemia progression within thymus vicinity might be predicted as blood circulating vasculature has very limited participation in normal thymus development [13]. In contrast, VEGFR2-negative and lymphatic vessels were most closely associated with early leukemia dissemination, although they most likely serve as a scaffold. For example, we consistently observed that single Rag2+ cells emigrating from the thymus towards the kidney marrow appeared to embed/line up along the outer surface of fli1+/VEGFR2-vasculature. Although no active movement of Rag2+ cells occurred during our 30 minutes of observation (data not shown), we hypothesize that the cells slowly crawl along fli1+/VEGFR2-vasculature, gradually widening the migration trail at 21dpf, and later forming secondary infiltrates in the kidney area before disseminating throughout the body. We are in process of completing time-lapse imaging of up to 8 hours to better define how this migration occurs. Rag2:GFP-Myc leukemia cells also collectively migrated towards the dorsal-lateral part of the thymus, i.e. towards medulla, where they again lacked VEGFR2+ vascular supply but were closely associated with fli1+/VEGFR2- and lymphatic vessels, occasionally intravasating into the latter.

Interestingly, we also consistently observed that a subpopulation of Rag2:GFP-Myc leukemic cells broke the basal membrane in the ventral part of the thymus and spread towards oxygen-rich aortic arches. This phenomenon demonstrates a spatiotemporal uncoupling of leukemia dissemination and invasion within the same tumor, leading us to hypothesize that leukemic cell fate and intra-tumoral heterogeneity may be influenced by vascular context, in which the metabolic, chemokine and cellular composition of the blood vessels, microvasculature, and lymphatics within areas of the thymus differ and can influence leukemia behavior. For example, normal thymocytes in the ventral cortical area of thymus display the highest rate of proliferation during development due to proximity to the oxygen–rich pharyngeal epithelial region [35], and it is possible that leukemic cells that we observed migrating towards oxygen-rich areas of VEGFR-positive vasculature were similarly in cell cycle. Additionally, differential availability of oxygen depending on the vascular composition of the niche may create regional differences in metabolism that favors one behavior in leukemic cells over another. For example, the proximity to VEGFR2+ vessels that carry oxygen-rich erythrocytes would increase the oxygen gradient in the tissue, which favors oxidative phosphorylation, while erythrocyte depleted microvasculature (fli1+; VEGFR2-) and lymphatics (lyve1+) would generate microenvironment that would favor a more “hypoxic” metabolism in leukemia cells. We are currently investigating the specific mechanisms that control metabolic re-programming in leukemic cells in the primary site of leukemia development.

In total, considering the high level of consistency in migratory behavior of Rag2:GFP-Myc expressing leukemia across individual leukemia-bearing zebrafish, the acquisition of random mutations that can collaborate with Myc expression to drive leukemia onset and progression seems very unlikely. We hypothesize that the microenvironment-derived metabolic, growth factor, and cytokine cues create a cascade of intrinsic and migratory changes in leukemia cells that lead to leukemia spread in the initial stages of the disease, and this may be regulated in large part by specific vascular networks. While vascular networks are described in great detail in early embryonic development in zebrafish (up to 5dpf), they have not been explored at the same levels in juvenile and adult animals; this investigation is warranted and has increasing significance since the majority of pathological processes, cancer in particular, typically occur in later stages of ontogenesis with high specification of vascular networks in relation to tissue/organ type. Visualization of these interactions will greatly enhance our understanding of systemic interactions between cancer cells and their microenvironment, and will establish novel and more specific targets for antitumor therapy.

## 4. Materials and Methods

### 4.1 Zebrafish Husbandry

All experimental procedures involving zebrafish were approved by the University of Kentucky’s Institutional Animal Care and Use Committee, protocol number 2015-2225. Transgenic lines used in this study are shown in Table 1.

**Table 1.**
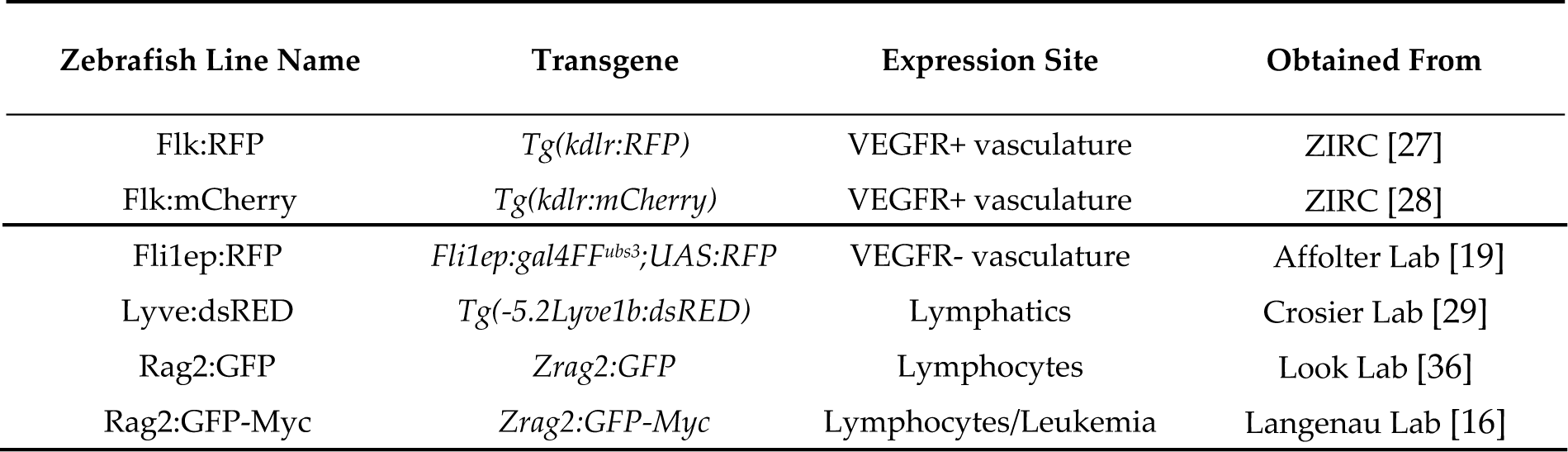
Zebrafish lines

### 4.2 Light Sheet Fluorescence Microscopy

To immobilize the fish for imaging, individual fish were transferred to 2mL microcentrifuge tubes, and excess water removed. 200μL of the E3 tricaine solution consisting of 2% of 4mg/mL Tricane (MS-222, Western Chemical Inc) with E3 media was added to the tubes, and fish were kept at rest for 5 minutes. 300μL of a solution of 3% SeqPlaque low-melt agarose (Lonza), melted in E3 media, was added to the tubes for a final agarose concentration of 1.5%. Tubes were mixed and the fish were loaded tail first into glass capillaries using a plunger (Zeis), and left for five minutes while the medium solidified. Agarose constricted fish were imaged using a Zeiss Light Sheet Z.1 Dual Illumination Microscope System and Zen imaging software. Fish aged 21 dpf or younger were imaged with a 20x objective lens and fish older than 21dpf were imaged with a 5x objective lens. Individual animals were used for every time point examined; the same animal was rarely used for sequential time points

### 4.3. Image Processing and Data Analysis

The images were analyzed using Imaris 9.2 (Bitplane). Files were converted to .IMS files using Imaris File Converter 9.3.0. The Imaris surface creation software was used enhance the clarity of the rag2:GFP-Myc and rag2:GFP cells in the images. This software automatically applies an easily visualized mask over all cells in the GFP channel. Images were saved as .tifs and Gimp photo editing software was used to deconvolute and crop images.

To quantify proximity of vasculature to the thymus, the average distance between the center of the thymus and the four largest surrounding vessels was defined in Imaris 9.2. The center of the thymus was determined by creating a cubic region of interest around the edge of the thymus and finding the midpoint of each dimension of the cube. The four largest vessels in the viewing field were determined by diameter. Spots were placed on these vessels where they closest to the thymus, and the Imaris 3D distance measurement feature was used to record distance of thymic center to vessel. Data represent the average of the four vessel distances for each image that was quantified.

Statistical significance was calculated using unpaired, two-tailed t-tests in GraphPad Prism. Significance was assigned when p<0.05.

## Author Contributions

The project was conceptualized by S.R. and J.S.B.; methodology developed by S.R; data collected and analyzed by S.R., M.B., S.P., E.H; schematics created by S.P.; writing—original draft preparation, S.R. and M.B.; writing—review and editing, S.R. and J.S.B; funding acquired by J.S.B.

## Abbreviations

dpf: Days Post Fertilization
GFP: Green Fluorescent Protein
Rag2: Recombination Activating Gene 2
RFP: Red Fluorescent Protein
VEGFA: Vascular Endothelial Growth Factor A
VEGFR2: Vascular Endothelial Growth Factor Receptor 2

